# Shaping the mechanical properties of a gelatin hydrogel interface via amination

**DOI:** 10.1101/2024.10.14.618299

**Authors:** Génesis Ríos Adorno, Kyle B. Timmer, Raul A. Sun Han Chang, Jiachun Shi, Simon A. Rogers, Brendan A. C. Harley

**Author notes:** **Corresponding Author:** B.A.C. Harley, Dept. of Chemical and Biomolecular Engineering, Cancer Center at Illinois, Carl R. Woese Institute for Genomic Biology, University of Illinois at Urbana-Champaign, 110 Roger Adams Laboratory, 600 S. Mathews Ave., Urbana, IL 61801, United States.

## Abstract

Injuries to musculoskeletal interfaces, such as the tendon-to-bone insertion of the rotator cuff, present significant physiological and clinical challenges for repair due to complex gradients of structure, composition, and cellularity. Advances in interface tissue engineering require stratified biomaterials able to both provide local instructive signals to support multiple tissue phenotypes while also reducing the risk of strain concentrations and failure at the transition between dissimilar materials. Here, we describe adaptation of a thiolated gelatin (Gel-SH) hydrogel via selective amination of carboxylic acid subunits on the gelatin backbone. The magnitude and kinetics of HRP-mediated primary crosslinking and carbodiimide-mediated secondary crosslinking reactions can be tuned through amination and thiolation of carboxylic acid subunits on the gelatin backbone. We also show that a stratified biomaterial comprised of mineralized (bone-mimetic) and non-mineralized (tendon-mimetic) collagen scaffold compartments linked by an aminated Gel-SH hydrogel demonstrate improved mechanical performance and reduced strain concentrations. Together, these results highlight significant mechanical advantages that can be derived from modifying the gelatin macromer via controlled amination and thiolation and suggest an avenue for tuning the mechanical performance of hydrogel interfaces within stratified biomaterials.

## 1. Introduction

The tendon-to-bone junction, or enthesis, is a transitional tissue that connects tendon and bone via a fibrocartilage interface [1]. Rotator cuff tears are common enthesis injuries, representing 4.5 million patients and 7 billion US dollars spent annually on treatments in the United States alone [2]. While rehabilitative treatments are options for partial injuries, full thickness tears typically require surgical intervention, leading to approximately 250,000 rotator cuff surgeries performed annually in the United States [3]. Such repairs are clinically challenging, with larger tears suffering re-failure rates as high as 94% [4–6]. While failures can be attributed to a variety of complications, current surgical techniques often fail to promote functional regeneration of the tendon-to-bone enthesis [1]. The most common surgical treatment for rotator cuff repair is mechanical fixation of the torn tendon to the underlying bone, and results in the formation of functionally inferior fibrous scar. While the native fibrocartilaginous enthesis provides a compliant tissue interface able to dissipate local strains [7–10], the fibrous scar does not provide a similar mechanical advantage, making it more difficult to dissipate forces and leading to new tears [11]. Because of these shortcomings as well as increasing need for rotator cuff treatment due to our active and ageing population, alternative strategies that prioritize enthesis regeneration are urgently needed.

We have previously reported a stratified biomaterial scaffold for use in rotator cuff repair. We developed lyophilization approaches to integrate porous tendon- (structurally anisotropic) and bone-mimetic (calcium phosphate mineralized, structurally isotropic) collagen scaffolds that can induce local tenogenic or osteogenic mesenchymal stem cell activity without the need for differentiation media or exogenous factor supplementation [12–16]. While biphasic (tendon-bone) scaffolds displayed significant strain concentrations at the point of integration, we recently added a compliant thiolated polyethylene glycol (PEG-SH) hydrogel zone between scaffold compartments to reduce strain concentration and failure significantly [17]. This method used enzymatic crosslinking of a thiolated PEG macromer, allowing an interfacial hydrogel to crosslink while diffusing into the adjacent collagen precursor suspensions prior to lyophilization. This process resulted in a continuous hydrogel transitional zone rather than three discrete material zones. However, while synthetic materials like PEG offer mechanical advantages, they do not naturally promote cellular adhesion or motifs appropriate for remodeling, making them less suitable for regenerative biomaterials. This shortcoming inspired our investigation of gelatin, a denatured form of collagen, as a base material for a second-generation triphasic scaffold. Gelatin contains cell attachment (RGD) and remodeling (e.g., metalloproteinase, MMP) motifs, and it can be functionalized for a variety of crosslinking approaches, including the thiol-based crosslinking [18–20]. We have shown thiolated gelatin hydrogels (Gel-SH) support mesenchymal stem cell (MSC) chondrogenic differentiation as well as the emergence of signatures of fibrocartilage differentiation in response to paracrine signals generated by MSCs seeded in tendon- and bone-mimetic collagen scaffolds [21, 22]. These inspire a Gel-SH insertional zone in our triphasic scaffold.

Gel-SH hydrogel macromers are formed via insertion of thiol groups onto the gelatin backbone but are limited by the concentration of amine groups amenable to thiolation. Duggan et al. described a promising method for increased gelatin thiolation by substituting carboxylic acid groups on the native gelatin backbone for additional free amines using ethylenediamine and EDAC (1-ethyl-3-(3-dimethylaminopropyl) carbodiimide) [18]. While carbodiimide-based crosslinking of primary free amines and carboxylic acids can be employed after a hydrogel gelation process to improve mechanical wet properties [23–25], increasing the overall extent of gelatin thiolation suggests a strategy to tune mechanical performance, thought at a potential cost of reduced access to secondary carbodiimide crosslinking. Here, we define the impact of tuning the degree of gelatin amination during the fabrication of a Gel-SH macromer, and consider the balance between primary (dithiol-based) and secondary (carbodiimide-based) crosslinking on hydrogel stability, mechanical properties, and use as a hydrogel transition in an interfacial biomaterial. We hypothesize that increasing the degree of gelatin amination (shifting the balance to increased dithiol primary crosslinking) will increase mechanical performance of the monolithic hydrogel as well as triphasic biomaterial containing a Gel-SH hydrogel insertion.

## 2. Materials and Methods

### 2.1. Amination of gelatin (Gel-NH_2_)

Type B, bovine-sourced gelatin (Sigma Aldrich, St. Louis, MO, 225 Bloom) was dissolved in half reaction volume (reaction volume: 50mL per gram of gelatin) of phosphate buffer (pH 5.0, 0.1 M) at 50 °C. Ethylenediamine (Thermo Fisher Scientific, Waltham, MA) of the desired mass ratio to gelatin (0.35, 1.05, or 2.8 g per 1 g gelatin) was then added [18]. The pH of the reaction was adjusted to 5 using 10 N HCl (Fisher Scientific, Hampton, NH), with temperature kept below 50 °C using ice packs to avoid gelatin denaturation. Once the reaction reached room temperature, EDAC (1-ethyl-3-(3-dimethylaminopropyl) carbodiimide hydrochloride, Oakwood Chemical, Estill, SC) was added in a 5.6:1 mass ratio (ethylenediamine: EDAC) [18]. The solution was then brought to total reaction volume with phosphate buffer and allowed to react for 24 hours. Following this, aminated gelatin was dialyzed (cellulose based membrane, MWCO: 12-14 kDa) for 5 days in deionized water (DI H_2_O), frozen at −20 °C, and lyophilized (FreeZone 2.5L Benchtop Freeze Dryer, Labconco Corp., Kansas City, MO) for 7 days.

### 2.2. Quantification of primary free amines in gelatin backbone

Primary free amines on aminated gelatin were quantified via 2,4,6-Trinitrobenzenesulfonic acid (TNBS, Sigma-Aldrich, St. Louis, MO) assay [26]. Briefly, samples were dissolved in a sodium bicarbonate buffer (0.1M NaHCO_3_, pH 8.5) at 1 mg/mL at 40 °C for 30 minutes. L-alanine (Sigma-Aldrich, St. Louis, MO) was dissolved in buffer and used as standard (0 – 3 mM). Next, 1 mL of 0.1 w/v% TNBS was added to each sample and standard, reacting at 40 °C for 2 hours. Finally, 1 N HCl (1 mL) and 10% SDS (0.5 mL, Sodium dodecyl sulfate, Sigma-Aldrich, St. Louis, MO) were added to each sample and standard. The absorbance of the samples was measured at 335 nm using a Tecan Infinite M200 Plate Reader (Tecan Männerdorf, Switzerland). Amine quantification was performed before and after gelatin amination to determine carboxylic acid conversion to primary amine, assuming 1:1 conversion. The starting concentration of carboxylic acid on the gelatin macromer was taken as 1000-1150 µmoles/g, as previously reported [18, 27].

### 2.3. Thiolation of gelatin (Gel-SH)

Gelatin thiolation was adapted from previous outlined methods [18, 21]. Native or aminated gelatin was dissolved in DI H_2_O (100 mL per 1 g gelatin) at 50 °C. Traut’s reagent (Thermo Fisher Scientific, Waltham, MA) was added at a 2:1 molar ratio relative to free amine concentration. The pH of the solution was adjusted to 7 with 10 N NaOH (VWR, West Chester, PA) for 20 minutes, then adjusted to pH 5 with 10 N HCl and reacted for 24 hours. Thiolated gelatin was then dialyzed (cellulose based membrane, MWCO: 12-14 kDa) in 5 mM HCl for 24 hours, dialyzed in 1 mM HCl for an additional 24 hours, frozen at −20 °C, and lyophilized (FreeZone 2.5L Benchtop Freeze Dryer, Labconco Corp., Kansas City, MO) for 7 days.

### 2.4 Quantification of free thiols in gelatin backbone

Free thiols on thiolated gelatin were quantified via Ellman’s assay (5,5-dithio-bis-(2-nitrobenzoic acid) [28]. Briefly, thiolated gelatin samples (5 mg/mL), L-cysteine standards (0 – 1.5 mM, Sigma-Aldrich, St. Louis, MO), and 4-5 mg/mL of Ellman’s reagent (Thermo Fisher Scientific, Waltham, MA) were dissolved in a sodium phosphate buffer (0.1M NaH_2_PO_4_, 1 mM EDTA, pH 8) at 40°C. Samples and standards (125 µL) were mixed with 1250 µL of buffer and 25 µL of Ellman’s reagent solution, vortexed, and allowed to react for exactly 15 minutes at 35 °C. Sample absorbance at 412 nm was measured in triplicate using a Tecan Infinite M200 Plate Reader (Tecan, Männerdorf, Switzerland).

### 2.5. Gel-SH hydrogel primary (1°) crosslinking

Gel-SH hydrogels were fabricated using an HRP-catalyzed radical reaction to form disulfide bond crosslinking [29]. The crosslinking reaction was performed at final concentrations of 5 mM tyramine (99% Tyramine, Sigma-Aldrich, St. Louis, MO), 10 mM H_2_O_2_ (30% Hydrogen Peroxide solution, Macron, Radnor, PA), and 5 U/mL Horseradish peroxidase (HRP, Thermo Fisher Scientific, Waltham, MA). Gel-SH was dissolved in Dulbecco’s phosphate buffered saline (DPBS, pH 7.3, Corning cellgro, Corning, New York) at 10 w/v % and 47 °C. During this process, a reaction solution of tyramine and H_2_O_2_ was prepared in DPBS. Upon complete Gel-SH dissolution in DPBS, HRP was added to the reaction solution and incubated 1 minute at room temperature to allow for activation. Stock solution of Gel-SH and reaction solution were then combined in appropriate amounts to yield a solution of 3.5 w/v % Gel-SH in solution with the crosslinking reaction reagents at specified levels.

### 2.6. Gel-SH primary (1°) crosslinking rheological characterization

Small amplitude oscillatory shear (SAOS) rheology was conducted to characterize the Gel-SH system linear viscoelastic properties. For each sample, 2 mL Gel-SH precursor solution was prepared as described above and immediately loaded onto the bottom plate (50 mm, sandblasted) of a MCR 302 rheometer (Anton Paar, Graz, AUST) with a Peltier system controlling the temperature at 25 ± 0.1 °C. A parallel plate fixture (50mm diameter, sandblasted) was used with a measuring gap of 500 μm. Oscillatory shear was continuously applied at 0.8 rad/s in the linear viscoelastic (LVE) regime (at a strain amplitude of 2%) to monitor gelation at a sampling interval of 7.9 seconds. For frequency sweep measurements, 1 mL hydrogel samples were prepared, loaded into a cylindrical polytetrafluoroethylene (PTFE) molds (diameter 25 mm, height 1 mm), and cured for 24 hours to fully crosslink before linear-regime frequency sweeps (at strain amplitude of 2%) were applied across a range of angular frequencies from 0.1 rad/s to 100 rad/s.

An amplitude sweep from 0.01 – 100% strain at 0.8 rad/s was performed to confirm LVE region and yield stress properties.

### 2.7. Gel-SH secondary (2°) crosslinking and rheological characterization

Gel-SH hydrogels were prepared as described, loaded into PTFE molds (diameter 25 mm, height 1 mm) and gelled for 1 hour in the presence of enzymatic agent HRP. Hydrogels were transferred into DPBS solutions for 23 hours to finish the crosslinking process at room temperature. Following this, a secondary crosslinking process was performed via exposure to 1-ethyl-3-(3-dimethylaminopropyl) carbodiimide hydrochloride (EDAC, Sigma-Aldrich) and N-hydroxysulfosuccinimide (NHS, Sigma Aldrich) at room temperature, as described previously [25, 30]. Briefly, hydrogels were transferred into PBS for 1 hour, secondary crosslinking was performed by subsequently soaking hydrated Gel-SH hydrogels for 90 minutes in a molar ratio (5:2:1) of EDAC and NHS relative to carboxyl groups in the scaffold (calculated as a function of scaffold mass, based on native gelatin molecular weight and composition). As EDAC:NHS are catalysts for crosslinking, hydrogels were then washed in PBS 3 times for 30 minutes to remove remaining EDAC:NHS. Rheological characterization was then performed (frequency and amplitude sweeps) as described in the previous section.

### 2.8. Sterilization, hydration, and secondary crosslinking of freeze-dried Gel-SH hydrogels

#### 2.8.1. Lyophilization of primary crosslinked Gel-SH hydrogels

Immediately following mixing, 100 µL of Gel-SH hydrogel precursor solution (3.5 w/v% Gel-SH, 5 mM tyramine, 10 mM H_2_O_2_ and 5 U/mL HRP) was pipetted into cylindrical PTFE molds (diameter 5 mm, height 5 mm) and allowed to crosslink for 1 hour. Following gelation, Gel-SH hydrogels were removed from the wells, placed on an aluminum plate, and lyophilized in a Genesis 25XL freeze-dryer (VirTis). Gels were first cooled to −10 °C at a rate of −1 °C min^-1^ and held at that temperature for 2 hours; following this, gels were brought up to 0 °C at a similar rate and sublimated at 0.2 Torr.

#### 2.8.2. Sterilization, hydration and secondary crosslinking

Lyophilized Gel-SH hydrogels were sterilized via ethylene oxide treatment on a 12 hour cycle using an AN74i Anprolene gas sterilizer (Andersen Sterilizers Inc, Haw River, NC), as in previous studies [31, 32]. Freeze-dried hydrogels were hydrated and crosslinked with EDAC:NHS at room temperature, as described previously [25, 30]. Briefly, hydrogels were soaked in 100% Ethanol for 2 hours, followed by multiple washes in PBS. Secondary crosslinking was achieved by soaking hydrated Gel-SH for 90 minutes in a molar ratio (5:2:1) of EDAC and NHS relative to carboxyl groups in the scaffold. Primary crosslinked constructs (lacking EDAC:NHS exposure) were created as controls by soaking in PBS at room temperature during this step. Following secondary crosslinking, hydrogels were washed multiple times with PBS to remove EDAC:NHS and stored in PBS for 24 hours prior to use. For materials prepared for cell culture, hydrogels were hydrated for 48 hours in mesenchymal stem cell growth media (low glucose Dulbecco’s Modified Eagle Medium with added glutamine (School of Chemical Sciences Cell Media Facility, University of Illinois Urbana-Champaign), 10% mesenchymal stem cell fetal bovine serum (Gemini Bio Products, West Sacramento, CA), and 1% antibiotic-antimycotic (Gibco, Billings, MA)) at 37 °C 5% CO_2_ prior to cell seeding.

### 2.9. Equilibrium water content (EWC) of freeze-dried Gel-SH hydrogels

EWC of Gel-SH hydrogels was conducted as previously described [29]. 100 µL freeze-dried Gel-SH hydrogels (n=6) were weighed to determine dried weight *(m_D_)* and soaked in 1 mL of DPBS for 24 hours for swollen mass after hydrating *(m_h_).* EWC% was defined as 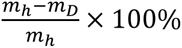 [33].

### 2.10. Mesenchymal stem cell culture on Gel-SH hydrogels

Human mesenchymal stem cells derived from bone-morrow (hMSCs, RoosterBio, MD) were cultured in RoosterNourish-MSC (RoosterBio, MD) at 37 °C and 5% CO_2_ and expanded to passage 5. hMSCs were seeded in Gel-SH hydrogel following a static point seeding method at 150,000 cells/hydrogel, based on previously established protocols [22]. Briefly, a 5 µL cell solution of appropriate concentration was incubated on top of each hydrogel for 2 hours to allow for cellular infiltration into the construct. Following this, 1 mL of mesenchymal stem cell growth media was added to each hydrogel. Complete growth media was changed every 3 days until experimental completion.

### 2.11. Cell bioactivity assessment

Metabolic activity of hMSCs was measured at experimental Days 1 and 7 using an alamarBlue assay (n=6, Invitrogen, Carlsbad, CA) [16, 34]. Hydrogels were washed with PBS (phosphate buffered saline) and incubated with 9:1 volume ratio of complete growth media: alamarBlue for 90 minutes at 37 °C, 5% CO_2_ and 100 rpm. Hydrogels were then returned to normal growth media, and aliquots of 100 µL incubated solution were measured in triplicate for fluorescence at 540/580 nm using a F200 spectrophotometer (Tecan, Männedorf, Switzerland). A standard curve at experimental Day 0 was generated to correlate fluorescent reading with approximate cell count.

Cell number was directly assessed as a function of total isolated DNA at experimental Days 1 and 7 using a Zymo Quick-DNA Microprep Plus Kit (n=5, Zymo Research Crop, Irvine, CA). A standard curve at experimental Day 0 was generated using known quantities of hMSCs and following the “Biological Fluids & Cells” protocol as directed, with all samples centrifuged at 15,000g at relevant steps and eluted into 10 µL. DNA was isolated from hMSCs within Gel-SH hydrogels by following the “Solid Tissues” kit protocol as directed, with tissue samples incubated in Solid Tissue Buffer and Proteinase K at 55 °C overnight and spun for all relevant steps at 15,000g. All isolated DNA was quantified in duplicate via a Nanodrop Lite spectrophotometer (ThermoFisher).

Cells were assessed visually via live-cell fluorescent imaging after 9 days of culture. Cell-laden hydrogel samples were washed with PBS for 5 minutes and stained with a live/dead staining kit containing calcein-AM / ethidium homodimer-1 (L3223, ThermoFisher) according to the manufacturer-directed protocol. Fluorescent imaging was captured on a DMi8 confocal microscope (Leica) at either 488 or 561 nm for live and dead imaging, respectively. Images colorization and contrast were adjusted in ImageJ for clearer viewing of cells.

### 2.12. Collagen-GAG suspension

#### 2.12.1. Mineralized collagen suspension

Mineralized collagen suspension was fabricated according to previously established methods [35–37]. Briefly, type I bovine Collagen (1.9 w/v%, Collagen Matrix Inc., NJ), chondroitin-6-sulfate from bovine cartilage (0.84 w/v%, CAS: 9082-07-09, Spectrum Chemical Mfg. Corp., CA), and calcium salts (0.9 w/v% CaOH and 0.4 w/v.% Ca(NO_3_)_2_•4H_2_O, Sigma-Aldrich, St. Louis, MO) were homogenized in 0.1456 M phosphoric acid/0.037 M CaOH mineralized buffer. Suspension was stored at 4 °C and degassed prior use, as previously described.

#### 2.12.2. Non-mineralized collagen suspension

Non-mineralized collagen suspension was fabricated as described before [38, 39]. Briefly, type I bovine Collagen (0.5 w/v%) and chondroitin-6-sulfate (0.044 w/v%), were homogenized in 0.05M acetic acid. The suspension was stored at 4 °C and degassed prior use.

### 2.13. Triphasic scaffold fabrication

Triphasic collagen-GAG scaffolds were fabricated based on a previously established process [17]. A simplified PTFE mold with a ¼-inch thick copper side was used to contain liquid collagen suspensions and the crosslinking Gel-SH hydrogel during the diffusive and lyophilization processes, and a 3D-printed polylactic acid (PLA) phase divider was used to aid in the initial loading of different suspensions. Each mold well (30 mm long, 14 mm wide, 6 mm deep) was filled with the following components: 1050 µL non-mineralized collagen suspension directly against copper side of the well, 1050 µL mineralized collagen suspension on the opposite end of the well, and 450 µL of Gel-SH precursor in between. Immediately following the loading of all three components, the divider was removed, and the three solutions were allowed to integrate at room temperature for 1 hour. Following this period, the suspension-loaded mold was placed into a freeze-dryer and lyophilized under the same conditions described for monolithic Gel-SH hydrogels.

### 2.14. Triphasic uniaxial tensile testing

#### 2.14.1 Embedding and coating

In preparation for mechanical testing, triphasic scaffolds were halved (width: 7 mm each) and cut to uniform length (28 mm), ensuring equal distance between regions across samples. Scaffold ends were then embedded into custom 3D-printed PLA endblocks via a two-component silicone potting and encapsulating compound (RTV615, Momentive Specialty Chemicals Inc., Columbus, OH), with a modified ratio of 4:1 Component A: Component B, as previously established [17]. The compound was allowed to partially cure (approximately 20 minutes at 60 °C) and then coated upon the endblock inside surfaces. Scaffold ends were inserted into the coated endblocks, placed onto glass slides, and maintained at 60 °C overnight to ensure complete fixation. Scaffolds were prepared for digital image correlation (DIC) [40] via speckled-patterning with waterproof India ink (BLICK Art Materials, Galesburg, IL) using a gravity feed airbrush (Master Airbrush, U.S. Art Supply) fitted with a 0.3 mm nozzle set piece [17]. Scaffolds were allowed to dry overnight prior to DIC.

#### 2.14.2 Uniaxial tensile testing

Uniaxial tensile testing was performed using an Instron 5943 Testing System (Instron, Norwood, MA, United States) with a 5-N electromechanical load cell. Scaffolds were held in place using pneumatic grips (BioPuls Submersible Pneumatic Side Action Tensile Grips, Instron, Norwood, MA) clamped to the endblocks. Gauge length and cross-sectional area were measured for each sample, and scaffolds were strained at a rate of 1 mm min^-1^ until fracture [41]. From the stress-strain curve data generated, Young’s tensile modulus and toughness were calculated as the slope of the linear region of the curve and the area under the curve, respectively [17, 25]. Ultimate tensile stress (the maximum tensile stress recorded over the measurement period) and yield strain (the strain at which the scaffold either definitively failed, resulting in an end of measurement, or experienced significant, unrecoverable deformation) were also recorded.

### 2.15. Local strain mapping using digital image correlation

As in previous work, embedded and speckled patterned scaffolds were imaged and processed via digital image correlation (DIC) for local strain mapping at corresponding bulk scaffold strain rates [17]. During uniaxial tensing, scaffolds were imaged using a Canon EOS 5DS R DLSR camera fitted with a Canon Macro 100-mm lens (Canon, Tokyo, Japan) and illuminated with a SugarCube LED Illumination System set to an intensity of one (USHIO, Tokyo, Japan). A timelapse remote was used to capture images at a consistent frequency (1 image per 6 seconds) throughout the course of uniaxial tensile testing and scaffold fracture. Images were labeled with the measured bulk strain corresponding to the time of capture and correlated using a variation of the MATLAB file package “Digital Image Correlation and Tracking” (Copyright ©2010, C. Eberl, D.S. Gianola, and S. Bundschuh) modified by E. Jones (Improved Digital Image Correlation version 4 – Copyright © 2013, 2014, 2015 by E. Jones).

In running DIC correlations, the region of interest was set to the entire scaffold, with a step size of 20 pixels (1200-1500 pixels total). Image correlations were conducted with a subset size of 41 pixels, a threshold of 0.5, and a search zone of 2. Correlation displacements were smoothed and calculated into strain values based on scaffold gauge length (approximately 13µm / pixel), with a smoothening kernel size and number of smoothing passes set to 11 and 3, respectively, and a cubic finite element displacement interpolation. Localized strain (E_yy_) was then visualized as a heat map at various bulk strain rates. Finally, local strain data was compared between samples by exporting data as an average of line scans. For each sample, vertical line scans (from tendon to bone) were taken at 3 x-proportions (0.25, 0.5, 0.75), and the average of these values was plotted as the average local strain as function of position in mm.

### 2.16. Statistics and data visualization

All statistical analyses were run through R (4.3.2) via RStudio (2023.12.1+402). Prior to evaluation, raw data sets were screened for outliers using a Grubbs outlier test. Any determined outliers were removed for statistical testing, though they remain visible in any visualized data. Data sets were then fitted to a linear model and tested for normality (Shapiro-Wilk) and equal variance (Levene), with a p-value of 0.05 needed to reject the null hypothesis. For independent data with normal distribution and equal variance, an ANOVA test and Tukey’s post-hoc test were used to determine significance (p < 0.05). Data with equal variance but a non-normal distribution was evaluated by Kruskal Wallis Dunn (Benjamini-Hochberg) tests, and data with normal distribution but unequal variance was tested by Welch’s ANOVA and a Games-Howell post-hoc. Finally, for data neither a normal distribution nor equal variance between groups, significance was determined via Welch’s Heteroscedastic F test with trimmed means and winsorized variances and a Games-Howell post-hoc test.

Figures were created with OriginPro. Box plot whiskers and diamond plot points denote ± 1.5 interquartile range, with a straight line and a point indicating the median and mean, respectively. In bar plots, the data is represented as the average ± 1 standard deviation, with straight lines denoting the median. Groups with different letter in data plots denotes statistical significance (p <0.05)

## 3. Results

### 3.1. Degree of functionalization accelerates Gel-SH gelation process

We fabricated a series of Gel-SH variants with increasing degree of R-SH substitution via the addition of increasing amounts of ethylenediamine (0 g, 0.35 g, 1.05 g, 2.8 g) per gram of gelatin. TNBS and Ellman’s assays were used to characterize the concentrations of primary free amines and thiols substitutions, respectively, on the gelatin backbone. Broadly, we found that increased concentrations of free amines on the gelatin backbone resulted in greater degrees of thiolation (Figure 2a). Characterization via small amplitude oscillatory shear (SAOS) rheology suggested that an increased degree of amination accelerated Gel-SH crosslinking, though long periods of time (> 8 hours) were necessary to reach equilibrium in all samples (Figure 2b). We characterized key kinetics parameters in this process: the *critical time* (t_crit_) at which the gelation process starts to be measurable (10% of the maximum rate) [29]; the *gelation time* (t_gel_) between the start and end of measurable changes in the storage modulus, G’; and the time at which the rate of change of G’ (dG’/dt) reaches its maximum (t_max_) (Figure 2c, S1a). Variations in amination significantly impacted kinetics of initial gelation, with aminated variants (0.35, 1.05, 2.8 g) initiating gelation significantly earlier than the non-aminated (0 g) Gel-SH (p = 0.016, Figure 2d). Greater degree of amination also accelerated the overall crosslinking process, as noted by a significantly reduced t_gel_ value (p=0.019; Figure 2e, S1a-b).

**Figure 1.**
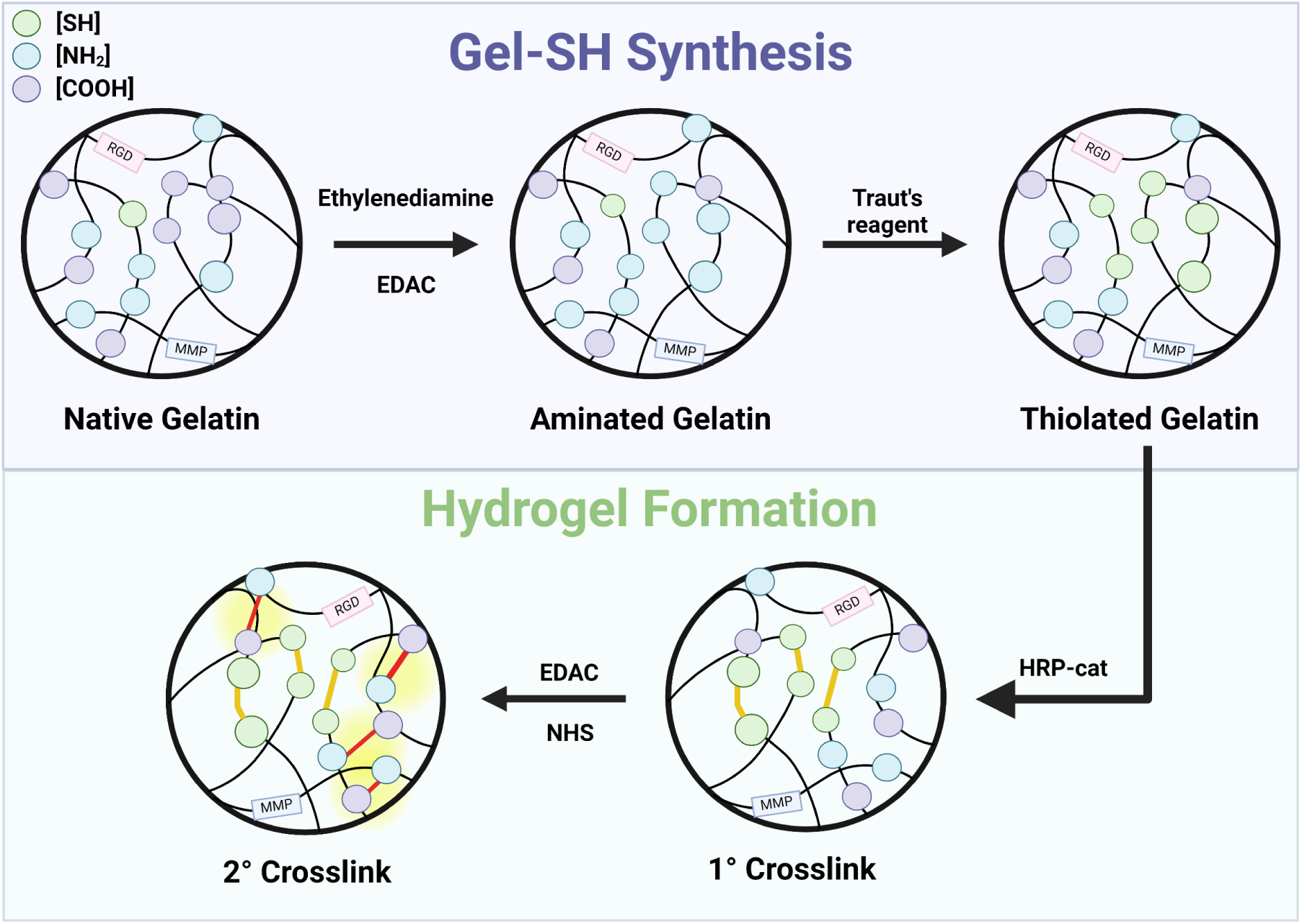
Gel-SH synthesis and hydrogel formation. Native gelatin is aminated by replacing carboxylic acid functional groups along the gelatin backbone with amines using ethylenediamine and EDAC. Traut’s reagent is then employed to substitute amines with thiols to create thiolated gelatin (Gel-SH). Crosslinked hydrogels are fabricated by oxidizing Gel-SH thiol groups to form disulfide bonds following an enzymatically (HRP) catalyzed reaction. Remaining primary amines and carboxylic acids are available to be crosslinked using carbodiimide (EDAC:NHS) crosslinking.

**Figure 2.**
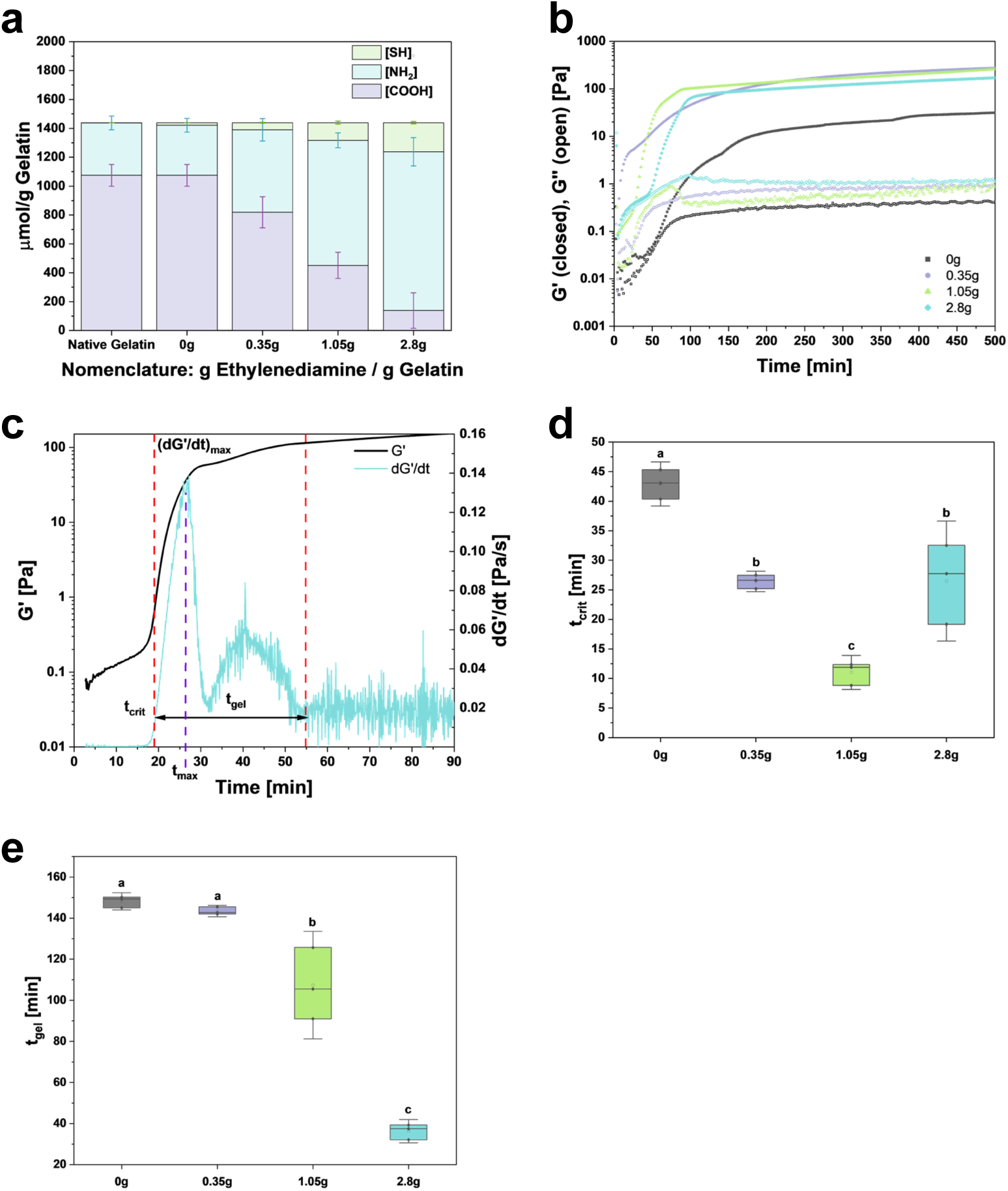
Primary crosslinked Gel-SH kinetic properties. **a.** Material functional group concentration: free thiols [SH], primary free amines [NH_2_], and carboxylic acids [COOH], as measured by Ellman’s, and TNBS assays. **b.** SAOS time sweep at an angular frequency of 0.8 rad/s and a strain amplitude of 2% used to determine kinetic parameters of Gel-SH hydrogels. **c.** Representative diagram of time sweep characterization for 2.8g. Values were evaluated by taking the first derivative of storage modulus with respect time, with t_crit_ being the time in which the slope of G’ change abruptly and the 10% of the maximum rate and t_gel_ is the width of the band. **d.** Calculated t_crit_ values for all Gel-SH types. **e.** Calculated t_gel_ values for all Gel-SH types. (n=3). Different letters denote statistical significance (p<0.05) between groups.

### 3.2. Secondary carbodiimide crosslinking alters rheological properties of modified Gel-SH

We subsequently evaluated the steady-state rheological properties for all Gel-SH variants both after primary crosslinking (thiol-mediated gelation) and after secondary crosslinking via EDAC:NHS carbodiimide crosslinking. Hydrogels were characterized after 24 hours of fabrication via amplitude and frequency sweeps to ensure complete full gelation from primary crosslinking. Amplitude sweeps at 0.8 rad/s confirmed all measurements were performed under the LVE region at 2% strain; all variants displayed overshoots in G’’ (the loss modulus), a characteristic behavior of yield stress materials (Figure 3a-d). Frequency sweeps were used to determine final material rheological properties; little to no time dependence was observed for Gel-SH variants after either primary or secondary crosslinking (Figure 3e-f, S2a-d). G’ at 0.1 rad/s from the frequency sweep, associated with long time scale deformation in the timeframe tested, was used to compare groups (Figure 3g). Gel-SH variants with higher degrees of amination were associated with a significantly reduced G’ relative to the unmodified 0 g Gel-SH (p = 0.025). Interestingly, higher degree of amination also significantly reduced the effect of secondary EDAC:NHS crosslinking on the G’, suggesting the loss of carboxyl groups due to amination decreased the availability for NH_2_-COOH crosslinks (Figure 3g). Finally, measuring the equilibrium water content percent (EWC%) revealed that higher degree of gelatin amination leads to lower equilibrium water content for both 1° and 2° crosslinked hydrogels, with 1° crosslinked gels displaying higher EWC% than hydrogels after 2° crosslinking (lower EWC% suggests a more densely crosslinked hydrogel, with increasing crosslinks limiting water absorption).

**Figure 3.**
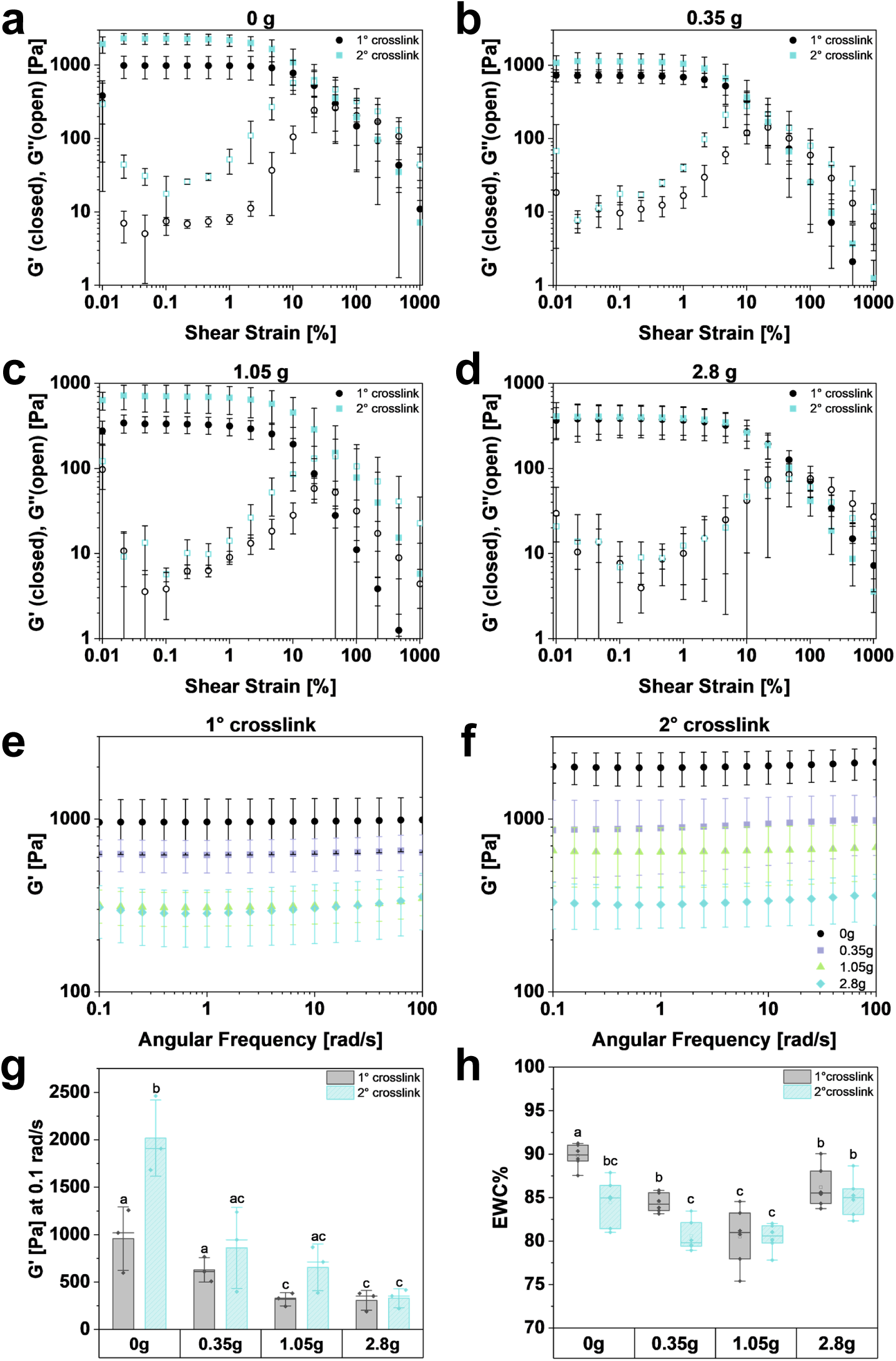
Secondary crosslinked Gel-SH mechanical effects. Amplitude sweeps at an angular frequency of 0.8 rad/s of primary and secondary crosslinked hydrogels of **a.** 0g, **b.** 0.35g, **c.** 1.05g and **d.** 2.8g. Frequency sweeps at a strain amplitude of 2% of **e.** primary and **f.** secondary crosslinked hydrogel variants. **g.** Storage modulus (G’) at equilibrium of Gel-SH variation after primary and secondary crosslinking (values at 0.1 rad/s, n=3). **h.** Equilibrium water content percentage (EWC%) of primary and secondary crosslinked Gel-SH variants (n=6). Different letters denote statistical differences (p<0.05) between groups.

### 3.3. Incorporation of modified Gel-SH alters global and local mechanical properties of triphasic collagen scaffolds

Following monolithic hydrogel characterization, we next evaluated the effect of incorporating Gel-SH with various degrees of amination within a multicompartment interfacial biomaterial. We measured mechanical properties of various triphasic scaffolds comprising a standardized non-mineralized collagen scaffold and mineralized collagen scaffold compartment linked by a Gel-SH interface. We observed that all Gel-SH variants successfully integrated between non-mineralized and mineralized collagen slurries to form stable triphasic scaffolds. Lyophilized triphasic scaffolds were tested under tensile deformation until fracture. Stress-strain curve was analyzed to identify several parameters, including the tensile Young’s modulus (defined by the linear elastic region), toughness (defined by the area under the stress-strain curve), ultimate tensile stress (maximum stress observed), and yield strain (strain at fracture) (Figure S3). We observed significant increases in the tensile Young’s modulus with increasing gelatin amination (2.8 g vs. 0.35 g; Figure 4a). Similar effects were observed with ultimate tensile stress (Figure 4c). We observed no significant differences in bulk toughness or yield strain between groups, though trends suggest a benefit with increased gelatin amination, particularly for the 1.05 g treatment (Figure 4b-c).

**Figure 4.**
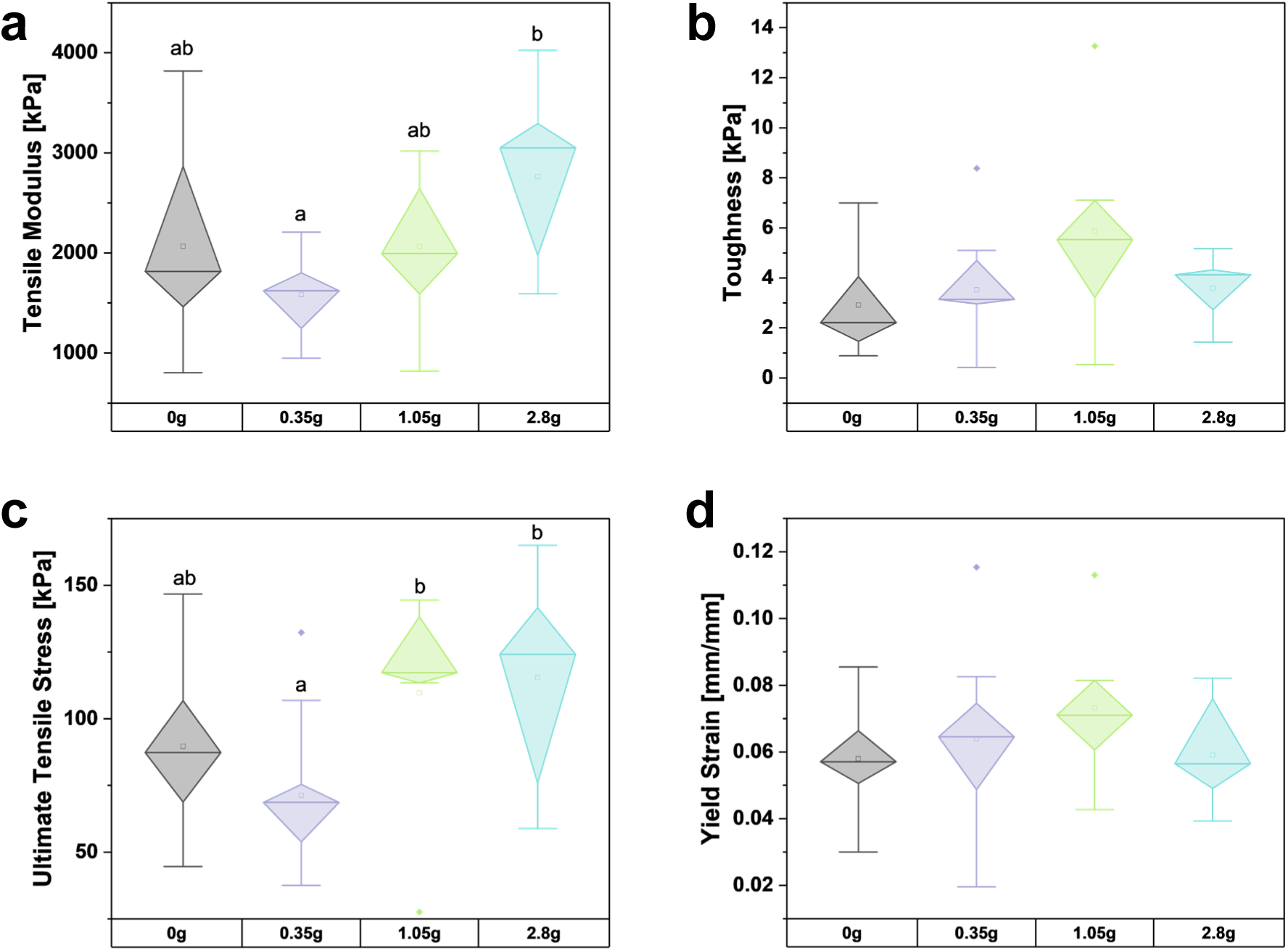
Bulk mechanical characteristics of triphasic collagen scaffolds containing a Gel-SH insertional zone. **a.** Tensile Young’s modulus (n = 15-16). **b.** Toughness (n = 8-11). **c.** Ultimate Tensile Stress (n = 8-11). **d.** Yield Strain (n = 8-11).

Following analysis of bulk material properties, we evaluated the local mechanical profile of the Gel-SH interface, quantifying local tensile strain in response to applied 2% bulk strain via DIC (Figure 5). Broadly, scaffolds containing low (0 g) or high (2.8 g) degrees of amination appeared to experience higher levels of local strain (>8%) nearer to the tendinous region of the scaffold as well as increased levels of localized strain (>2%) within the interfacial region itself. However, the incorporation of a Gel-SH interface using 0.35 g or 1.05 g Gel-SH variants resulted in lower maximum local strain (<7%) and generally lower levels of local strain within the hydrogel interface (Figure 5b-c). Representative DIC heat maps for each group are available in Figure S4a-d.

**Figure 5.**
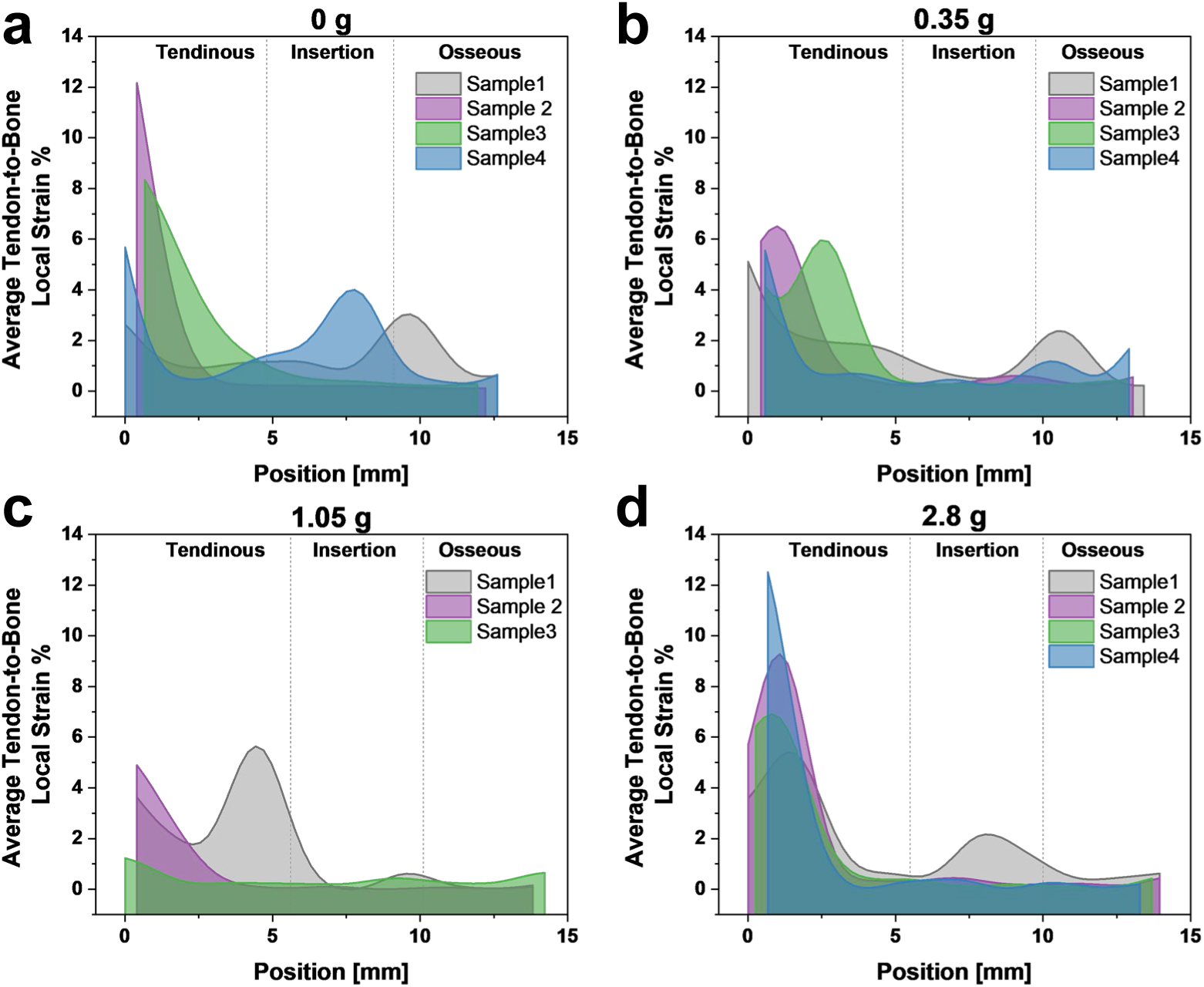
Local tensile strain profiles across triphasic collagen scaffolds containing a Gel-SH hydrogel insertion. Average local strain concentrations (n=3-4) as a function of distance along the length of the triphasic scaffold at ∼2% bulk strain for **a.** 0 g, **b.** 0.35 g, **c.** 1.05 g, and **d.** 2.8 g Gel-SH variants. Dotted lines indicate approximate boundaries between tendinous, insertion, and osseous regions.

### 3.4. Modified Gel-SH supports hMSC viability within the construct at levels equal to or greater than unmodified Gel-SH

Finally, we compared the bioactivity of seeded human mesenchymal stem cells (hMSCs) within Gel-SH variants by evaluating overall metabolic activity and resident cell number (Figure 6a-b) as well as live/dead imaging (Figure 6c). The 2.8 g Gel-SH group was not included in this analysis as it failed to rehydrate comparably with the other groups and did not significantly contribute to local strain reduction. hMSCs within the 0.35 g Gel-SH demonstrated greater bioactivity relative to the unmodified 0 g Gel-SH, as demonstrated by significantly greater metabolic activity as well as equivalent or greater cell proliferation without significant changes observed to cellular infiltration or morphology.

**Figure 6.**
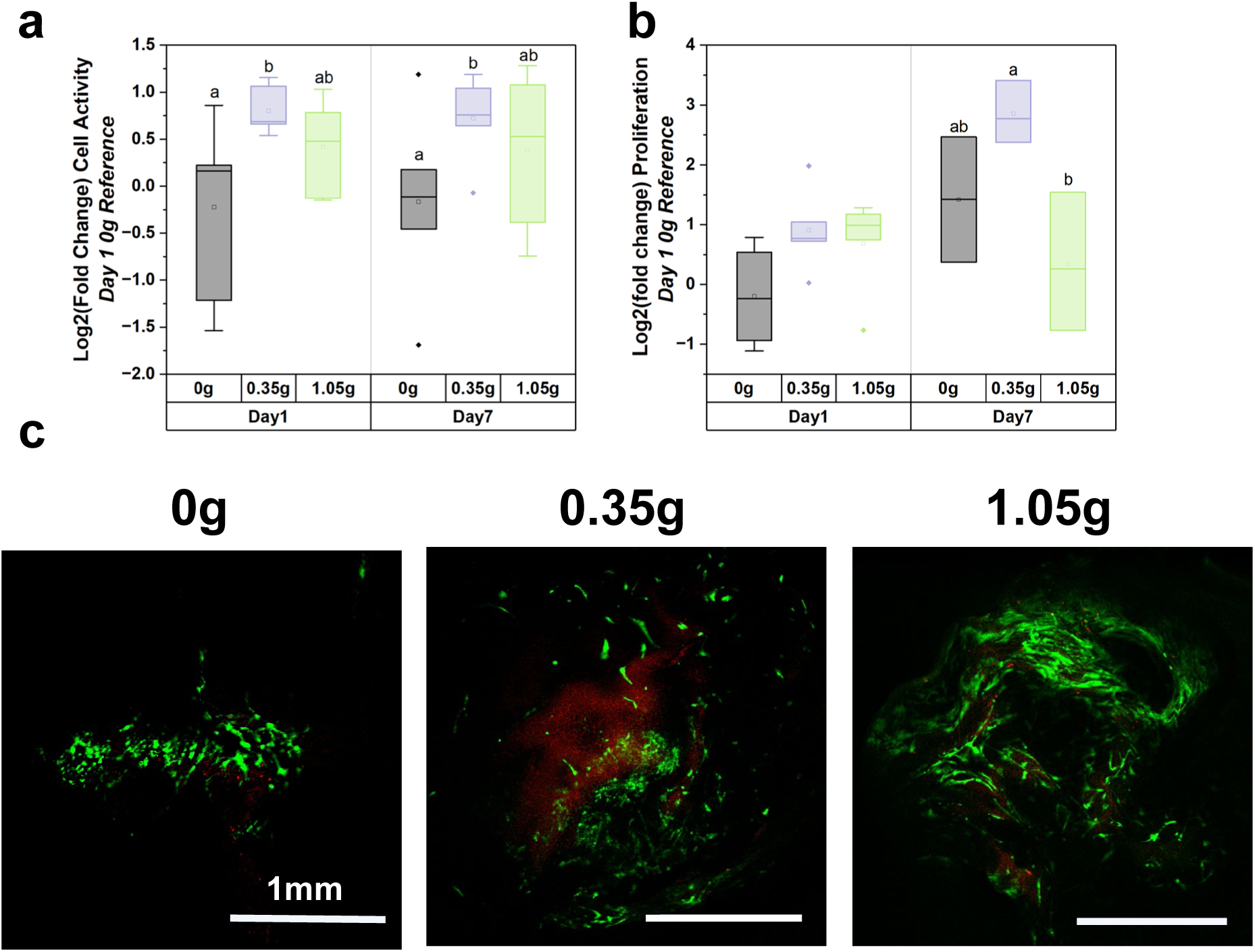
hMSC viability and proliferation on modified Gel-SH. **a.** Metabolic activity (n=6) after 1 vs. 7 days of culture on Gel-SH variants. **b.** Relative proliferation (n=3-5) as measured by DNA content after 1 vs. 7 days of culture. Relative values are normalized to the average value for the 0 g Gel-SH group at Day 1 and expressed as the log_2_(fold change). **c.** Representative fluorescent images of live-stained hMSCs within Gel-SH variants (Day 9).

## 4. Discussion

Naturally derived polymers, such as collagen and gelatin, offer significant advantages in biomaterial design due to their availability, presentation of native ligands, and capacity for degradation [42]. Scaffolds and hydrogels produced from gelatin and collagens have shown the capacity to support osteogenic and tenogenic differentiation of stem and progenitor cell populations, an essential quality in any biomaterial under development for tendon-to-bone enthesis repair [16, 43–46]. Previously, we described a multiphase material that, via incorporation of a thiolated polyethylene glycol (PEG-SH) hydrogel zone between dissimilar tendon and bone specific collagen scaffolds, was able to dissipate local interfacial strain that otherwise occur due to the sharp mechanical transition between dissimilar materials. In addition to the mechanical benefits of local strain dissipation, such changes could also reduce strain realized by seeded cells, improving viability. However, the PEG-SH interface did not support substantial cell viability and growth, limiting biological testing of this material [17]. Here, we explored modifications to a thiolated gelatin macromer, substituting carboxyl acid groups on the gelatin backbone with free amines and thiols to adjust the mechanical performance of the resultant crosslinked Gel-SH polymer network [47].

Thiol-mediated crosslinking is a powerful tool for enzymatic formation of stable, crosslinked hydrogel networks [18, 48]. While native gelatin contains few if any thiol groups, thiols can be installed via multiple mechanisms, notably by targeting amines on the gelatin backbone. However, thiolation is hindered by low concentration of primary amines on the gelatin backbone [18]. We hypothesized that improving the capacity of installing thiols on the gelatin backbone would provide a route to enhance gelatin mechanical properties within a stratified material. We first increased amine concentrations on the gelatin backbone to enable higher thiolation conversion, introducing amines on ethylenediamine then crosslinking them to carboxylic groups with EDAC. We generated multiple gelatin variants (labeled 0 g, 0.35 g, 1.05 g, 2.8 g, based on the ratio of grams of ethylenediamine per gram of gelatin) for subsequent analysis of rheological and mechanical performance. Our targeting of carboxyl groups on the gelatin backbone to enable amination and subsequent thiolation also introduced a secondary question. The triphasic (tendon-enthesis-bone) scaffold composite as well as the individual tendon and bone specific scaffolds are conventionally crosslinked via EDAC:NHS prior to use (to further increase scaffold mechanical performance) [39, 49]. However, the amination and then thiolation reactions reported here would alter the relative availability of the primary free amines and carboxylic acids sites on the gelatin backbone that are the targets for EDAC:NHS crosslinking [50]. Hence, tailoring the degree of amination and thiolation also introduced the opportunity to shape the relative importance of primary (thiol-mediated) versus secondary (amine/carboxylic acid mediated) crosslinking that could occur within the gelatin hydrogel insertion connecting dissimilar tendon and bone scaffolds.

Rheological and mechanical analysis demonstrated that gelatin amination prior to thiolation substantially modified the hydrogel gelation and steady state properties. Increasing the degree of amine substitution increased the number of installed thiol groups and accelerated metrics of gelation compared to non-modified (0 g) Gel-SH (Figure 2a-e). Interestingly, while amination variants displayed faster crosslinking kinetics, they also displayed lower average G’ values as well as a reduced effectiveness of secondary crosslinking (EDAC:NHS) for increasing steady-state elasticity. We had hypothesized that increased amination would increase bulk hydrogel stiffness by increasing thiol content and by extension primary crosslinking, but it is possible that high amounts of substituted thiols coupled with the floppy backbone of the gelatin macromer contributed to an increasing degree of self-crosslinks rather than networks between gelatin chains. Faster kinetics could also have contributed to a reduced potential to form more crosslinks where multiple available thiols are competing for a single binding site. A higher degree of modification would also reduce the number of available carboxylic acids sites for the secondary EDAC:NHS step, reducing the potential for COOH groups to be activated to form COOH-NH_2_ bonds and decreasing the efficacy of the secondary crosslinking of aminated gelatin. Broadly, this data highlights the potential for impactful functionalized group modification within Gel-SH, offering a wider range of gelatin macromer compositions amenable to different crosslinking techniques. Through these modifications, we can promote accelerated gelation to facilitate integration with collagen suspensions and a lyophilization process needed for biomaterials containing a continuous interface [41].

A major motivation for the modification of Gel-SH was the hypothesis that modifying the degree of amination could further reduce localized strain responses within our triphasic biomaterial scaffold design [41]. Strain concentrations at the interface between tendon and bone are a major contributor to the failure of surgical repairs to the rotator cuff, which fail to regenerate the mechanically-supportive tendon-to-bone junction and highlight the potential for a regenerative biomaterial that mimics this native function [7, 51]. Here, while modifications to the Gel-SH hydrogel interface did not significantly improve bulk mechanical properties (tensile modulus, toughness, etc.) of the triphasic biomaterial, we did observe improvement in local strain distributions when the triphasic material was under tensile load. Digital image correlation data showed that scaffolds fabricated with the 0.35 g or 1.05 g Gel-SH hydrogel variants at their interfaces experienced local strain concentrations that were less severe and less likely to occur within the insertion zone, instead occurring in more biologically preferable regions of flanking tendon and bone. Interestingly, while trends of increasing tensile modulus, ultimate tensile stress, and yield stress were observed with increasing degree of gelatin amination, these trends were not significant compared to the unmodified (0 g) group. This suggests that the amination process has more far-reaching effects on base gelatin and the end-product Gel-SH than simply increasing primary crosslinking, a conclusion that the compositional and rheological data would also support. This also suggests the need for future study of how shifts in gelation kinetics of the aminated Gel-SH macromer affect the degree of interdiffusion with the surrounding collagen suspension during lyophilization. We previously showed suspension freezing kinetics during lyophilization alters resultant pore microarchitecture [52, 53], so it is possible Gel-SH gelation and freezing kinetics contribute to this phenotype.

Finally, to ensure modifications and potential mechanical improvements did not require biological tradeoffs, we confirmed that all Gel-SH variants accommodated hMSC activity. Notably, rather than reducing cell viability, the 0.35 g and 1.05 g Gel-SH variants promoted significantly higher metabolic activity levels than the unmodified (0 g) group, with the 0.35 g Gel-SH variant promoting the greatest degree of hMSC proliferation. Ongoing work is considering the role that the degree of Gel-SH amination may have on the functional phenotype of seeded hMSCs; recently published work from our group showed that the secretome generated by hMSCs can promote aspects of enthesis-specific hMSC fibrochondrogenesis [22]. These findings set the stage for studies of hMSC activity in response to changes in monomer functional group composition within more complex environments, such as in the presence of cyclic tensile strain [49] or flanking tendon-and bone scaffold compartments [54].

## 5. Conclusions

We report an approach to create a library of thiolated gelatin (Gel-SH) materials via introduction of an intermediate amination step in the preparation of gelatin macromers. Altering the concentration of gelatin functional groups offers a pathway for tuning the kinetic and elastic features of the resultant hydrogel, addressing a key downside of naturally-derived hydrogel macromers. This work has demonstrated biological and mechanical improvement to a Gel-SH hydrogel under development for an enthesis tissue engineering application by sequential amination and thiolation of the gelatin backbone. The kinetics of the primary (thiol-mediated) crosslinking were faster in hydrogels with greater amination, though such modification also resulted in reduced deformation properties. Interestingly, when applied as an interface with a triphasic biomaterial linking dissimilar tendon- and bone-specific collagen scaffolds, Gel-SH variants with greater amination demonstrated more robust mechanical properties, with reduced local interfacial strain in response to tensile loading. These efforts suggest a path for expanded use of naturally derived biomaterial modifications and highlight opportunities to optimize hydrogel interface zone across multiple avenues of consideration (primary vs. secondary crosslinking; local strain performance) in the pursuit of interfacial biomaterials for regenerative repair of orthopedic insertions.

## Acknowledgements

The authors would like to acknowledge Michael Xu and Caleigh Arentsen (Materials Science and Engineering, Bioengineering, respectively; University of Illinois Urbana-Champaign) for assistance with data collection and analysis. The authors would also like to acknowledge the following institutes: the Carl R. Woese Institute for Genomic Biology and the Beckman Institute for Advanced Science and Technology. Research reported in this publication was supported by the National Institute of Arthritis and Musculoskeletal and Skin Diseases of the National Institutes of Health under Award Number R01 AR077858 (BACH) and the National Institute of Dental and Craniofacial Research of the National Institutes of Health under Award Number R01 DE030491 (BACH). We are grateful for the funding for this study provided by the NSF Graduate Research Fellowship DGE-1144245 (RSHC), the Grainger College of Engineering SURGE Fellowship (GRA) and Sloan UCEM Scholarship (GRA). The interpretations and conclusions presented are those of the authors and are not necessarily endorsed by the National Institutes of Health or the National Science Foundation.

## Contributions (CRediT: Contributor Roles Taxonomy [55, 56])

**G. Ríos Adorno:** Data curation, Formal Analysis, Visualization, Investigation, Methodology, Writing – original draft, Writing – review & editing. **K. Timmer:** Data curation, Formal Analysis, Visualization, Investigation, Methodology, Writing – original draft, Writing – review & editing. **R. Sun Han Chang:** Conceptualization, Data curation, Visualization, Investigation. **J. Shi:** Formal Analysis, Visualization, Investigation, Methodology. **S. Rogers:** Conceptualization, Methodology, Writing – review & editing. **Brendan Harley:** Conceptualization, Resources, Project administration, Funding acquisition, Supervision, Writing – review & editing.

