## Supplementary Materials for "Shaping the mechanical properties of a gelatin hydrogel interface via amination"

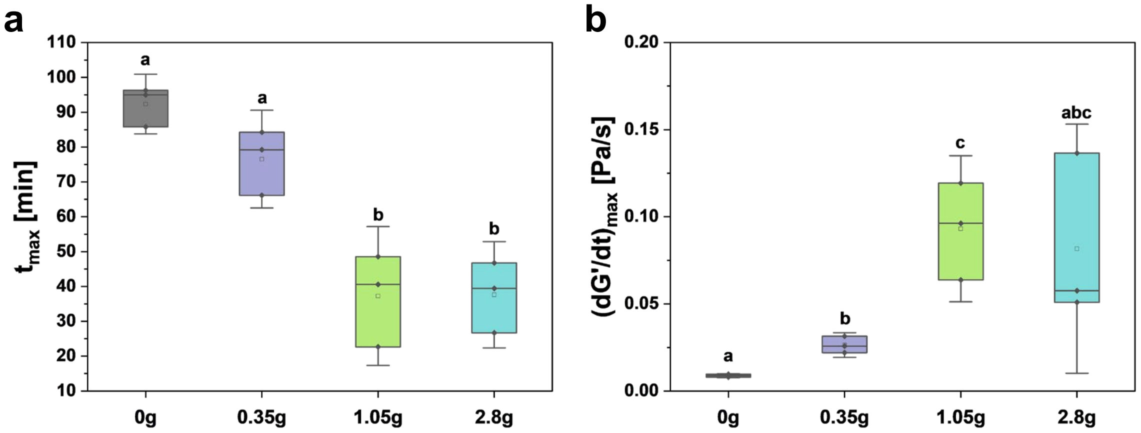


**Figure S1. Kinetics parameters for primary crosslinked Gel-SH. a**. Time at which the change of G' reaches its maximum, $t_{max}$. **b.** Maximum rate change of G', $(dG'/dt)_{max}$. Trends suggests right shifts on maximum peak for gelation, and increasing rate changes with higher degree of amination, indicative of faster gelation processes.

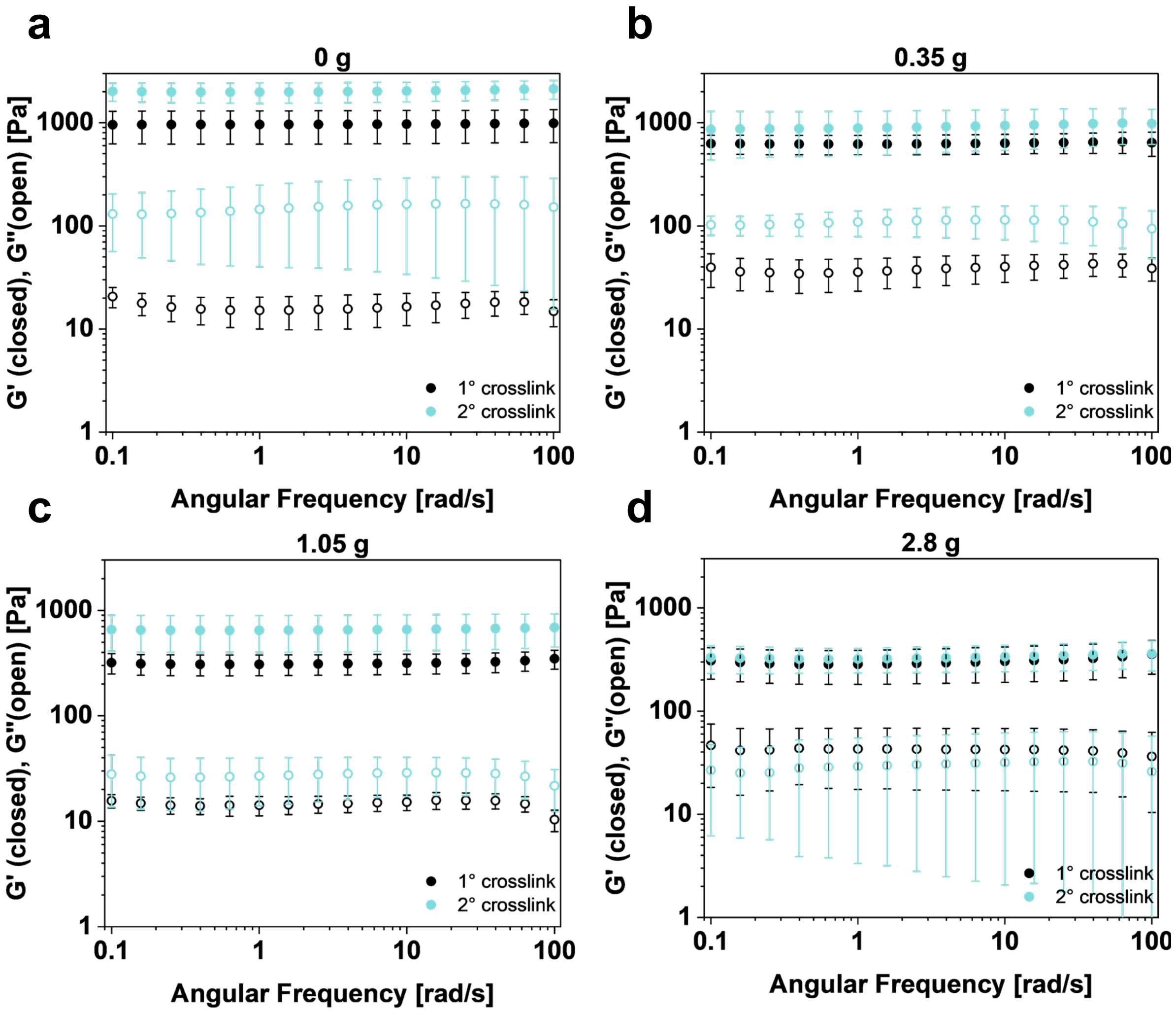


**Figure S2. Frequency Sweeps of primary and secondary crosslinked Gel-SH hydrogels at a strain amplitude of 2% of a.** 0 g, **b.** 0.35 g, **c.** 1.05 g, **d.** 2.8 g.

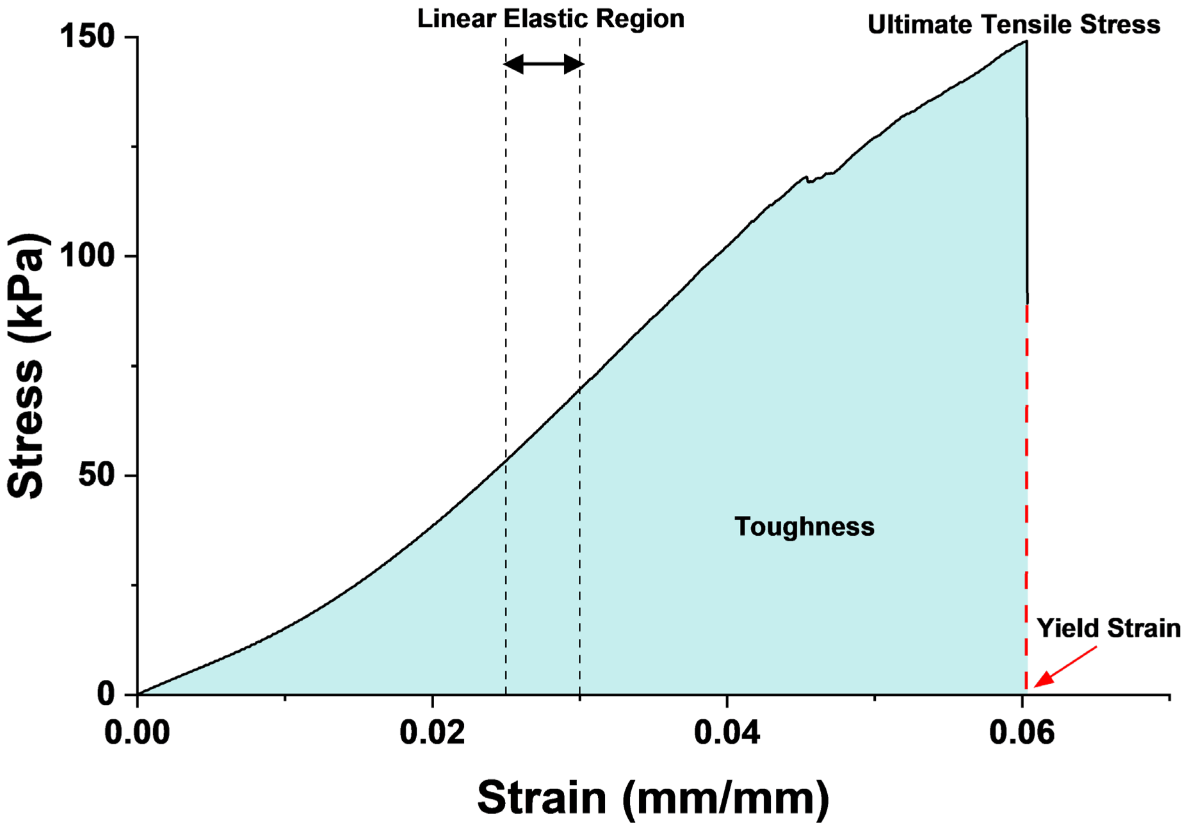


**Figure S3. Mechanical analysis of triphasic scaffolds under tensile deformation.** Stress-strain curves were obtained from tensile testing to calculate the tensile modulus, from the slope of the linear elastic region (linear regime 2.5% - 3% strain), toughness, the area under the stress-strain curve, the ultimate tensile stress, maximum experienced stress during the deformation and the yield strain, strain at which the material failed or experienced unrecoverable deformation.

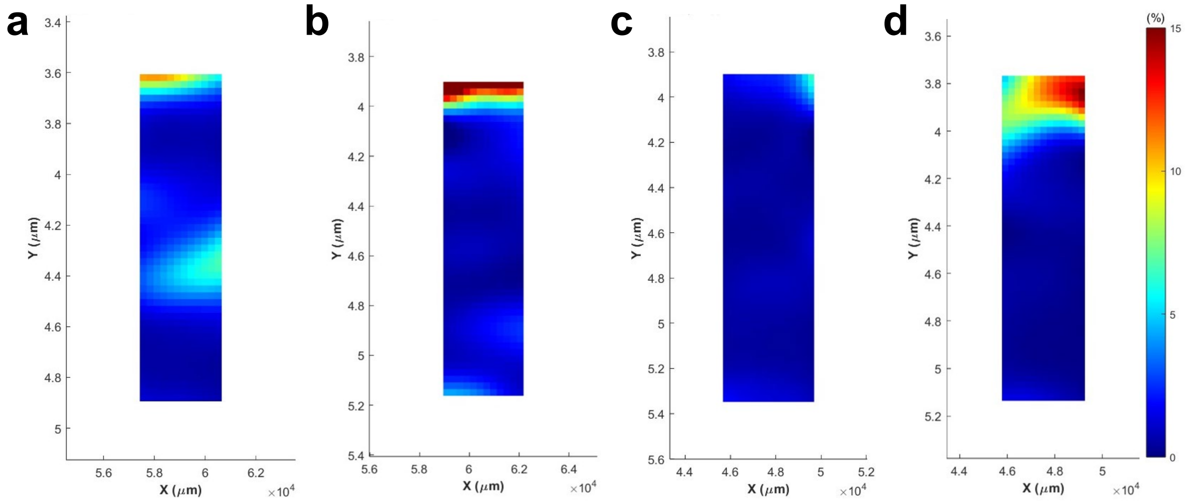


**Figure S4. Localized strain across the triphasic scaffold during tensile deformation.** Representative profiles of local strain at 2-3% applied strain with respect triphasic length of **a.** 0 g, **b.** 0.35g, **c.** 1.05 g and **d.** 2.8 g.
